# Responses to temperature shocks in *Zymoseptoria tritici* reveal specific transcriptional reprogramming and novel candidate genes for thermal adaptation

**DOI:** 10.1101/2024.12.16.628617

**Authors:** Silvia Minana-Posada, Alice Feurtey, Julien Alassimone, Bruce A. McDonald, Cécile Lorrain

**Affiliations:** Plant Pathology Group, Institute of Integrative Biology, ETH Zürich, Zürich, Switzerland

## Abstract

Pathogens’ responses to sudden temperature fluctuations, spanning various temporal scales, are critical determinants of their survival, growth, reproduction, and homeostasis. Here, we combined phenotyping, transcriptomics, and genome-wide association approaches to investigate how the wheat pathogen *Zymoseptoria tritici* responds to and recovers from temperature shocks. Survival emerged as the most significantly affected trait immediately following temperature shocks across 122 geographically diverse strains. In contrast, post-recovery phenotypic traits, including growth rate and melanization, showed no significant deviations from control conditions. Transcriptomic analyses of a reference strain revealed temperature stress-specific gene expression patterns, with genes involved in protein folding, redox homeostasis, membrane stabilization, and cell-wall remodeling playing central roles in the response. A multi-reference k-mer genome-wide association study (GWAS) identified six loci significantly associated with cold shock responses. Among these, two loci emerged as strong candidates for near-freezing temperature adaptation, including a 60S ribosomal protein gene involved in protein synthesis and stress recovery, and an NADH oxidoreductase gene implicated in redox homeostasis and oxidative stress tolerance. These findings shed light on the distinct molecular strategies *Z. tritici* employs to adapt to temperature stress and provide novel insights into fungal resilience under dynamic environmental conditions.

**Author summary:** Temperature fluctuations, an inherent aspect of natural environments, are increasingly exacerbated by climate change, intensifying challenges for organisms to maintain homeostasis amid more frequent and severe extreme weather events. This study reveals distinct phenotypic, transcriptomic, and genetic mechanisms underlying *Z. tritici*’s responses to short-term temperature shocks. Survival-related phenotypic traits were significantly reduced by heat and cold shocks, while other traits measured after a recovery period demonstrated the resilience of *Z. tritici* strains to temperature stress, reflecting efficient recovery mechanisms. Transcriptomic analyses uncovered temperature-specific gene expression patterns, emphasizing unique regulatory strategies, which mostly return to baseline levels after a recovery period. The discovery of novel loci associated with cold shock responses provides valuable insights into the genetic basis of resilience to short-term temperature stress, offering a foundation for future research on pathogen adaptation to fluctuating environments.

## Introduction

Fluctuations in temperature are an intrinsic feature of most ecological niches and can occur over different temporal scales—seasonally, diurnally, or abruptly during extreme weather events—posing challenges for organisms to maintain homeostasis and critical biological functions. The impact of such fluctuations has become increasingly pronounced in the context of global climate change, which has amplified the frequency, duration, and severity of extreme temperature events like heat waves and cold spells [1,2]. In many organisms, abrupt thermal shifts can significantly impact survival, growth, and reproduction, especially for ectothermic species like fungi. To cope with these challenges, fungi must employ rapid physiological and molecular responses to minimize cellular damage and maintain metabolic activity [3].

Understanding how fungal pathogens respond to sudden temperature changes is crucial, as their ability to adapt to fluctuating abiotic conditions directly impacts their survival, transmission, and infection severity [4–6]. These responses involve mechanisms such as protein stabilization, oxidative stress management, and membrane restructuring, which are essential for preserving cellular integrity during thermal shocks. Heat stress can trigger protein misfolding and cellular damage, leading to the activation of heat-shock proteins (HSPs) [7–9]. Cold stress disrupts metabolic activity, destabilizes RNA structures, and rigidifies membranes, often resulting in dehydration and ice-related damage [10–12]. Shared stress pathways, such as the high osmolarity glycerol (HOG) and cell-wall integrity (CWI) pathways [13,14], as well as reserve carbohydrates and protective pigments like melanin, also play pivotal roles in fungal resilience to temperature stresses [15,16]. These responses can vary significantly across fungal species and strains, influenced by genetic factors and prior exposure [17,18].

Fungi demonstrate exceptional thermal tolerance among eukaryotic organisms. Pathogens of endothermic hosts must survive in consistently high temperatures [19], whereas fungal pathogens of ectothermic hosts face variable thermal conditions, with seasonal and daily temperature shifts shaping their adaptation strategies [20,21]. For instance, human fungal pathogens such as *Aspergillus fumigatus*, can thrive at temperatures as high as 70°C and remain viable at −20°C [22,23]. Some fungal plant pathogens, such as the wheat pathogen *Zymoseptoria tritici* have evolved specific morphological structures associated with high-temperature stress responses [24,25]. Although the mechanisms involved in temperature stress response are well-characterized in model organisms such as yeasts, they remain understudied in most pathogenic fungi, particularly with regard to short-term temperature shocks [26]. Thus, combining phenotypic, genomic, and transcriptomic approaches offers unprecedented opportunities to dissect the genetic and molecular basis of fungal responses to thermal stress [27,28].

*Z. tritici,* the causal agent of Septoria Tritici Blotch (STB) in wheat, provides an ideal model for exploring fungal adaptation to heat and cold stress. As a fungal pathogen found in temperate regions, *Z. tritici* frequently encounters heat and cold stress, especially during seasonal transitions, requiring rapid physiological adjustments to maintain viability [29,30]. The global distribution of *Z. tritici* and extensive genomic resources, including 19 reference genomes spanning six continents, make *Z. tritici* ideal for linking genetic variation to phenotypic responses under fluctuating environmental conditions [31,32]. Recent studies have highlighted local adaptation in *Z. tritici*, with strains from warmer climates exhibiting elevated optimal temperatures [33,34]. Moreover, quantitative trait loci (QTL) mapping and genome-wide association studies (GWAS) have identified genetic regions associated with temperature adaptation [32,33,35–37]. However, the effects of abrupt temperature shocks remain largely unexplored.

In this study, we explored the phenotypic and transcriptomic responses of *Z. tritici* to short-term temperature shocks and examined the genetic basis underlying these responses. We hypothesized: (i) *Z. tritici* strains exhibit distinct phenotypic responses to short-term heat and cold shocks. These responses should show significant variation immediately after the shocks and after a recovery period, with different strains showing different responses; (ii) Heat and cold shocks elicit distinct gene expression profiles in *Z. tritici*, with each type of stress inducing a unique set of differentially expressed genes. We expect some overlap in the genes differentially expressed under both conditions; and, (iii) Heat and cold shock phenotypic responses are associated with distinct loci in *Z. tritici*.

## Results

### Heat and cold shocks impair the survival of *Z. tritici* spores

To investigate the impact of short-term temperature shocks on *Z. tritici*, we evaluated seven phenotypic variables in 122 strains from four geographically distinct populations (S1 Table) after 15-hour heat (30°C), cold (0°C), and control (18°C) treatments. Heat and cold stress treatments significantly impaired the spore concentration and survival rate (Fig 1A; S1 Fig; S2 Table; *p* < 0.05). The colony radius, melanization, melanization heterogeneity, growth rate, and the start of colony emergence traits were evaluated during subsequent growth at 18°C on PDA plates. While melanization exhibited a significant treatment effect when analyzed independently of timepoints, this significance disappeared when the interaction between treatment and timepoints (days post-inoculation) was included in the analysis. Similarly, colony radius, melanization heterogeneity, growth rate, and the timing of colony emergence did not show significant treatment effects, either as standalone factors or when considering their interaction with timepoints (Fig 1A; S1 Fig; S2 Table; p > 0.05).

**Fig 1.**
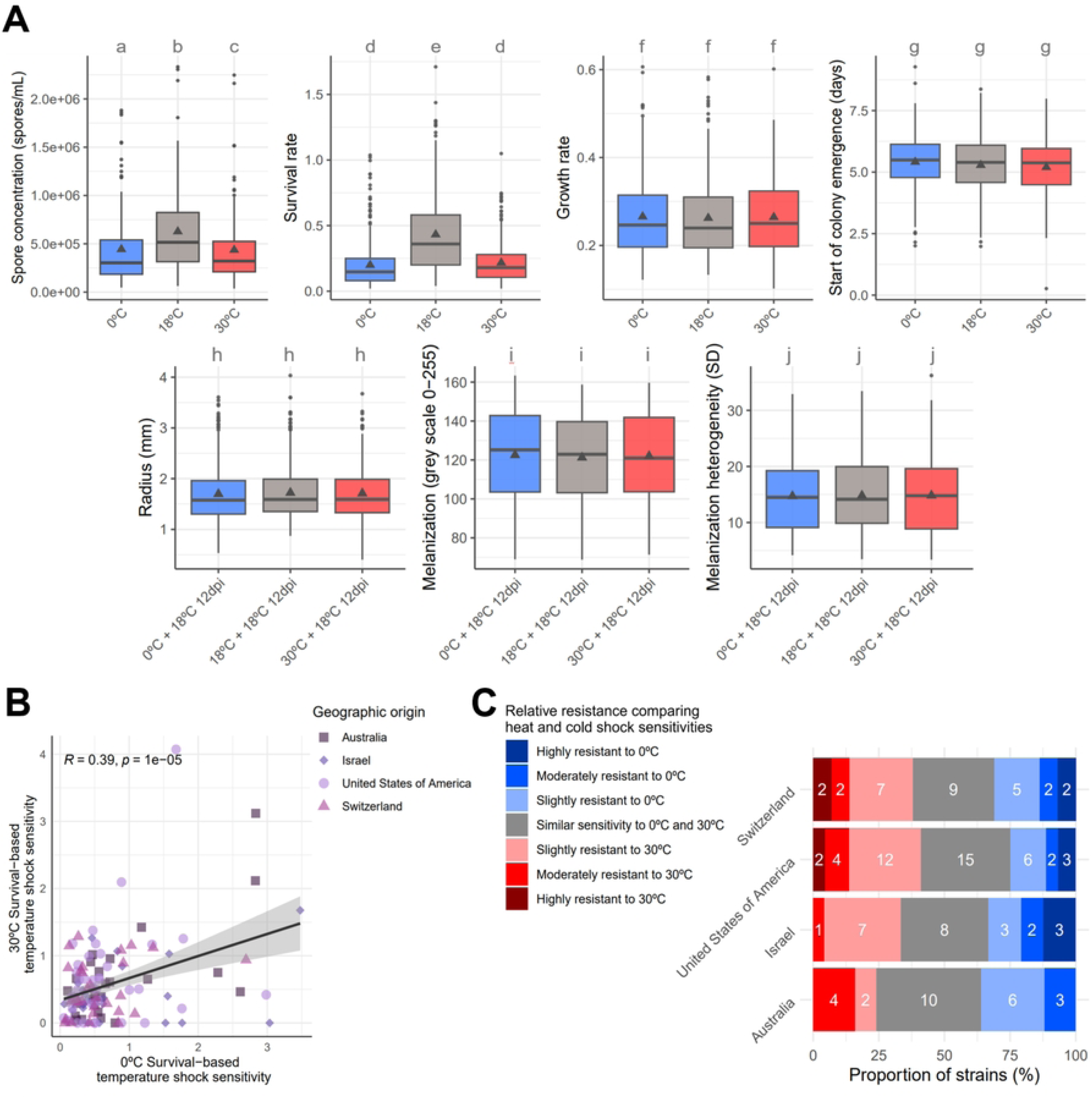
Treatment-specific and population-level responses in phenotypic traits and sensitivities to heat and cold shocks. (**A**) Boxplots displaying the seven phenotypic traits measured after temperature 15-hour temperature shock treatments. The triangle inside each box represents the mean. Treatments are as follows: 0°C for cold shock, 30°C for heat shock, and 18°C as the control. Lowercase letters (a–c) denote significant differences between treatments based on linear mixed-effects models per variable and Tukey’s post-hoc pairwise comparisons. *dpi*: days post-inoculation. (**B**) Scatter plot of survival-based sensitivity following heat shock (30°C) versus cold shock (0°C). Pearson’s correlation coefficient (*R*) and *p* are displayed at the top of the plot. (**C**) Proportions of *Zymoseptoria tritici* strains from each geographic population, categorized by relative resistance to temperature shocks. Categories are based on survival-based sensitivity ratios comparing heat shock (30°C) to cold shock (0°C). White numbers within the bars represent the total number of strains in each category (see S5-S6 Tables for details).

Both heat and cold shocks led to significantly lower spore concentrations compared to the control treatment (Fig 1A; S3 Table; Tukey’s post hoc test, *p* < 0.001). Spore concentrations were significantly lower after cold shock than after heat shock, indicating that 0°C caused greater impairment than 30°C (Fig 1A; S3 Table; Tukey’s post hoc test, *p* < 0.01). While spore concentrations were consistently lower after both temperature shocks across populations, no significant differences were observed between the heat and cold shock treatments within individual populations (S2 Fig; S3 Table). This reflects the considerable inter-strain variability in post-shock treatment spore concentration within *Z. tritici* populations. Similarly, heat and cold shocks significantly reduced survival rates (Fig 1A; S4 Table; Tukey’s post hoc test *p* < 0.001). However, survival rates after both shocks remained comparable, suggesting 30°C and 0°C imposed similar stress levels (Fig 1A; S4 Table; Tukey’s post hoc test, *p* > 0.05). This trend held across all populations (S3 Fig; S4 Table). These findings demonstrate that temperature shocks negatively impacted the spore concentration and survival rates of *Z. tritici* strains. However, once colonies emerged, they recovered rapidly, with no detectable effects on growth or morphology.

To examine potential cross-sensitivity between heat and cold shocks, we calculated survival-based sensitivity ratios for each treatment relative to the control (S4 Fig). A significant positive correlation was observed between heat and cold shock sensitivities (Pearson’s *R* = 0.39, *p* < 0.05; Fig 1B), suggesting that strains resistant to one stress were generally more resistant to the other. This correlation was significant in two of the four populations, while no significant relationship was detected for the Swiss and Israeli populations (S5 Fig). We classified strains into seven resistance categories to further explore strain-specific differences, comparing relative sensitivity to heat and cold shocks (S6 Fig; S5–S6 Tables). Most strains (42, 34%) exhibited similar sensitivity to both shocks, reflecting the observed cross-resistance. Smaller proportions displayed slight (28 strains, 23%) or moderate (11 strains, 9%) resistance to heat shock, while others showed slight (20 strains, 16%) or moderate (9 strains, 7%) resistance to cold shock. Few strains exhibited extreme specialization, with four strains (3%) highly resistant to heat shock and eight strains (7%) highly resistant to cold shock. These patterns appeared consistent across populations (Fig 1C). Overall, these results reveal a general pattern of cross-resistance between heat and cold shocks among *Z. tritici* strains, with most strains showing similar resistance to both conditions and only a minority demonstrating specialization.

### Distinct transcriptomic responses to heat and cold shock exposure

To investigate the gene expression response of *Z. tritici* to temperature shocks, we analyzed transcriptomic profiles of *Z. tritici* IPO323 spores immediately after 15 hours of temperature shock (*shock treatment*), as well as after an additional 75 hours at the control temperature (*shock-and-recovery treatment*). We performed a principal component analysis (PCA) using normalized read counts to assess the differences in transcriptomic profiles. The PCA revealed a clear separation between samples taken after the *shock treatment* and *shock-and-recovery treatmen*ts, suggesting that the primary variance captured reflects changes in the growth stage of the spores after 15h and 90h of growth, respectively (Fig 2A; PC1 explained 76% of the variance). Additionally, transcriptomic profiles differed between heat and cold *shock treatments* and the control (Fig 2A; PC2 explained 11% of the variance). However, after the recovery period, the transcriptomic profiles of both temperature treatments showed less differentiation from each other and the control (Fig 2A).

**Fig 2.**
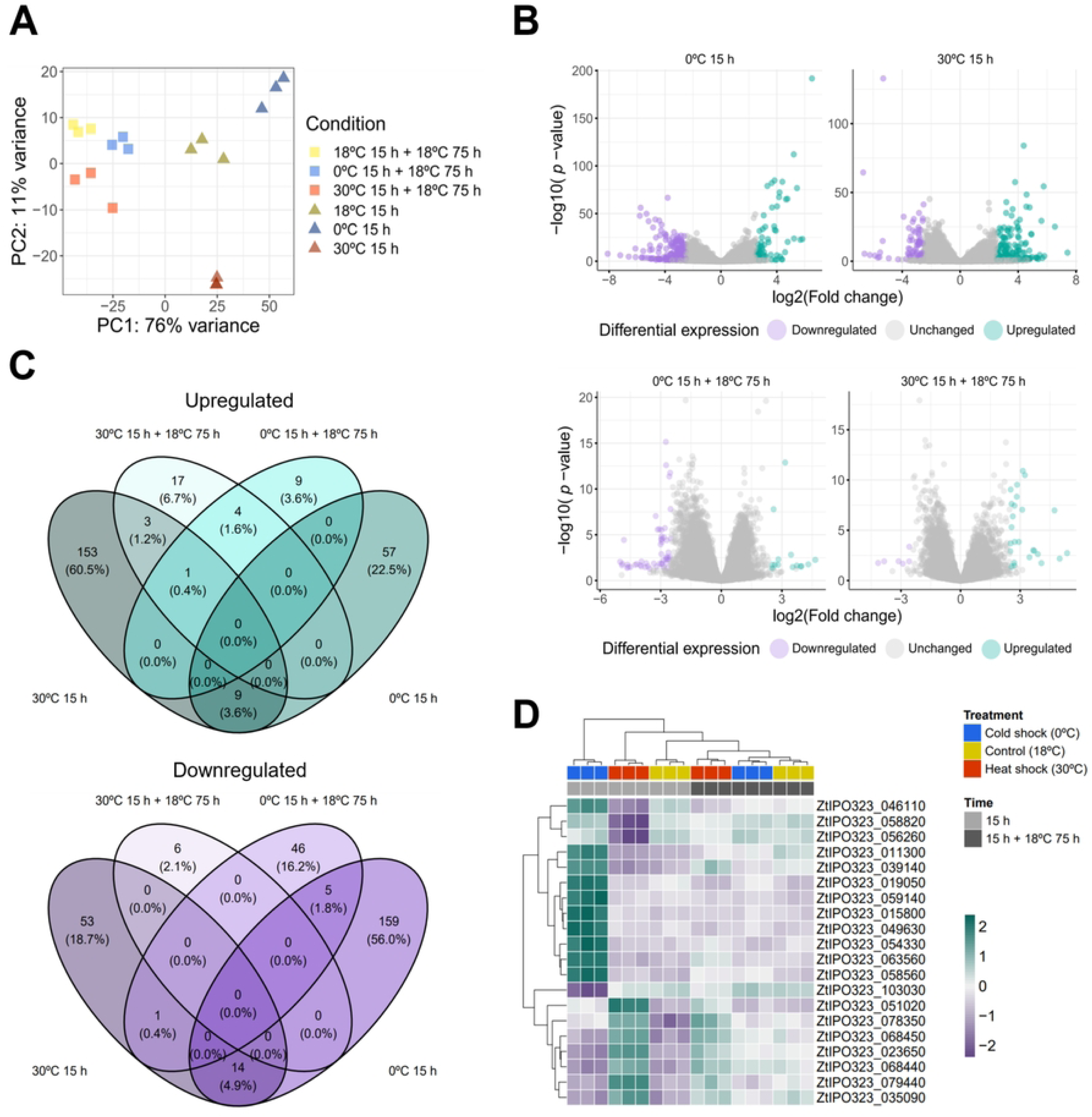
Transcriptomic analysis reveals distinct gene expression profiles in *Zymoseptoria tritici* following temperature shocks and recovery. (**A**) Principal component analysis (PCA) of transcriptomes from the reference strain IPO323, showing the first two principal components for samples after temperature shocks or control conditions at two timepoints. Triangles represent samples taken immediately after 15- hour temperature shocks, while squares indicate samples after an additional 75-hour recovery at the control temperature. Blue shapes denote cold shock (0°C), red denote heat shock (30°C), and yellow denote control (18°C). Each point represents a replicate. (**B**) Volcano plots illustrating differential gene expression (DEG) immediately after temperature shocks and following recovery. Genes with a log2-fold change > 2.5 are upregulated, while those < -2.5 are downregulated. (**C**) Venn diagrams portraying the overlap of upregulated and downregulated DEGs across temperature shocks and recovery conditions. (**D**) Heatmap of the 20 most significant DEGs (lowest adj. *p*), with columns representing replicates per treatment and rows representing each DEG.

We identified differentially expressed genes (DEGs) to further explore the effects of temperature shocks by comparing each shock treatment to the control (S7 Table). In total, we identified 449 DEGs (log2-fold change > 2.5 or < -2.5; adjusted *p* < 0.05) across the treatments. For heat shock, 153 genes were upregulated and 53 downregulated compared to the control (Fig 2B & C). In contrast, cold shock resulted in 57 upregulated and 159 downregulated genes (Fig 2B & C). The majority of DEGs after heat (65.8%) and cold (80.8%) shock treatments encode genes of unknown functions (S7 Fig). No gene ontology (GO) category was significantly enriched in any treatment. However, among the fraction of DEGs with predicted GO annotations, we found diverse functions, including “Carbohydrate transport and metabolism” (eight upregulated DEGs), “Secondary metabolites biosynthesis, transport and catabolism” (eight upregulated DEGs), “Amino acid transport and metabolism” (six upregulated DEGs) and “Posttranslational modification, protein turnover, chaperones” (four upregulated and five downregulated DEGs) in the heat shock treatment (S7 Fig). Similarly, we found functions including “Carbohydrate transport and metabolism” (five downregulated DEGs), “Secondary metabolites biosynthesis, transport and catabolism” (four downregulated DEGs), and “Amino acid transport and metabolism” (four downregulated DEGs) in the cold shock treatment (S7 Fig). We found similar functions differentially expressed in heat and cold shock treatments. However, these functions are overall upregulated in heat shock treatment while downregulated in cold shock treatment, strengthening the specific transcriptomic response to each temperature shock. We noted nine upregulated DEGs common to heat and cold shocks (Fig 2C). The common upregulated genes between heat and cold shock included a bZIP transcription factor-like gene (ZtIPO323_016070), a lipid transporter-like gene (ZtIPO323_069120), and seven genes of unknown functions that may contribute to general stress adaptation (S7 Table). Due to the established role of chaperone HSPs in temperature stress, we examined the expression of 25 predicted HSPs in *Z. tritici* strain IPO323 (S8 Fig; S7 Table). Cold shock led to the downregulation of HSP98 (ZtIPO323_029190), HSP20 (ZtIPO323_075280), and HSP30 (ZtIPO323_052640). Interestingly, HSP98 and HSP20 were also downregulated after heat shock. In contrast, HSP12 (ZtIPO323_063560) was upregulated under cold shock treatment. The remaining 21 HSPs showed no significant expression changes (S8 Fig; S7 Table). These findings suggest that specific HSPs contribute uniquely to cold and heat stress responses, with HSP12 potentially playing a central role in cold stress adaptation.

The shock-and-recovery treatments resulted in fewer DEGs, with 25 genes upregulated after heat recovery and 14 genes upregulated after cold recovery (Fig 2B & C). Among these, only four genes were upregulated in both treatments. In contrast, six genes were downregulated in the heat shock-and-recovery treatment, and 51 genes were downregulated in the cold shock-and-recovery treatment (S7 Table, S9 Fig). Few genes were differentially expressed in both the shock treatments and the shock-and-recovery treatments. For example, in the cold shock-and-recovery treatment, five downregulated DEGs, all uncharacterized proteins, were shared with the initial cold shock phase (S7 Table, S9 Fig). In the heat treatment, four overlapping upregulated DEGs between the shock and shock-and-recovery treatments included a Ca2+-modulated nonselective cation channel (ZtIPO323_041210), a small threonine-rich protein (ZtIPO323_091890), and two uncharacterized proteins (S7 Table).

We then focused on the 20 DEGs with the most significant adjusted *p* across all treatments. Of these, ten were associated with cold shock: nine upregulated and one downregulated (Figure 2D). The cold-responsive genes represented various functional categories, indicating a broad cellular response to cold stress. Among the upregulated genes, we found chaperones and stress-related genes such as: the HSP12 (ZtIPO323_063560), the stress-related RDS1 protein (ZtIPO323_039140), and a GroES- like chaperonin (ZtIPO323_054330). We also identified four genes related to metabolism and redox homeostasis, including an NAD(P)-binding protein (ZtIPO323_015800) and a NADPH-dependent medium-chain alcohol dehydrogenase (ZtIPO323_019050). The only downregulated DEG in this group was a NAD(P)-binding protein (ZtIPO323_103030). Additionally, we found two genes linked to transport and signaling, including an upregulated uracil permease-like protein (ZtIPO323_011300) and an upregulated AraC- like ligand-binding domain protein (ZtIPO323_049630). The remaining ten significant DEGs were linked to the heat shock treatment, with seven upregulated and three downregulated. Five upregulated and two downregulated genes were of unknown function. However, one upregulated DEG, predicted as a glycoside hydrolase family 128 protein (ZtIPO323_078350), might play a role in cell wall lysis and remodeling. We also identified two DEGs with signaling-related functions: one upregulated EF-hand calcium-binding protein (ZtIPO323_035090), and one downregulated aflatoxin regulatory protein (ZtIPO323_058820).

### K-mer GWAS identified a few loci associated with temperature shocks

We conducted k-mer-based genome-wide association studies (GWAS) across 122 *Z. tritici* strains to investigate the genetic basis of temperature stress responses. We analyzed seven phenotypic variables per treatment, resulting in 48 phenotypes. A total of 985 significant k-mers were identified across 26 phenotypes, using a 10% family-wise error rate threshold (S8 Table). Of these, 579 k-mers were associated with the control treatment, while 406 were specific to temperature shock conditions (S8 Table). Among the 406 k-mers linked to temperature shocks, 388 aligned with the 19 *Z. tritici* reference genomes. Further exploration of the genomic regions associated with these significant k- mers identified eight loci based on their proximity to orthologous genes (see Methods). After filtering out loci overlapping with transposable elements (TEs), six loci remained, each containing more than two significant k-mers (S9 Table). Notably, no loci associated with heat shock phenotypes were identified after this filtering, while six loci associated with cold shock phenotypes were found. These loci correspond to cold shock survival rate (1 locus), the start of colony emergence (2 loci), colony radius (2 loci), and melanization (1 locus). Three loci were shared across all 19 reference genomes, while three others were found in a subset of the reference genomes.

One locus on chromosome 12, containing between 14 and 90 significant k-mers, was associated with cold shock survival rate and was present in all 19 references (Fig 3A; S9 Table). This locus co-locates with one orthogroup (OG_2618) predicted to encode a NADH oxidoreductase 19.3 kDa subunit, part of the respiratory chain. For the start of colony emergence after cold shock, we identified a locus on chromosome 8, found in all 19 references (Fig 3B; S9 Table). This locus contained 3 to 22 significant k-mers and co- located with one orthogroup (OG_7328) encoding a 60S ribosomal protein gene. An additional locus associated with colony emergence was found on accessory chromosome 18 in references ST99CH_1E4 (17 significant k-mers), ST99CH_1A5, OregS90, and YEQ92 (3 significant k-mers). This locus falls within an intergenic region, with the nearest orthogroup (OG_12531) predicted to have an unknown function, located ∼900 bp downstream of the k-mer locus (Fig 3B; S9 Table). For colony radius after cold shock, we identified two loci. The first locus on chromosome 2, found in six references, contained nine significant k-mers (Fig 3C; S9 Table). The nearest orthogroup (OG_3769) is predicted to encode a transcription factor located ∼1.2 to 1.4 kb downstream of the significant k-mers. The second locus, found in 10 references on chromosome 11, contained three significant k-mers and overlapped with one orthogroup (OG_3167) encoding a chromate transporter domain-containing protein (Fig 3C; S9 Table). One significant locus associated with melanization was found on chromosome 3, containing ten significant k-mers that overlapped with one orthogroup (OG_4998) encoding a WD40/YVTN repeat-like-containing domain superfamily protein involved in protein binding (Fig 3D; S9 Table). A transposable element is annotated at the same locus in five references (ST99CH_3D1, ST99CH_3D7, I93, KE94, and OregS90) but not in the remaining 14 references. Hence, the k-mers GWAS identified a relatively small number of loci and genes specifically linked to temperature stress responses in *Z. tritici*, offering insights into genetic adaptation mechanisms across core and accessory chromosomes.

**Fig 3.**
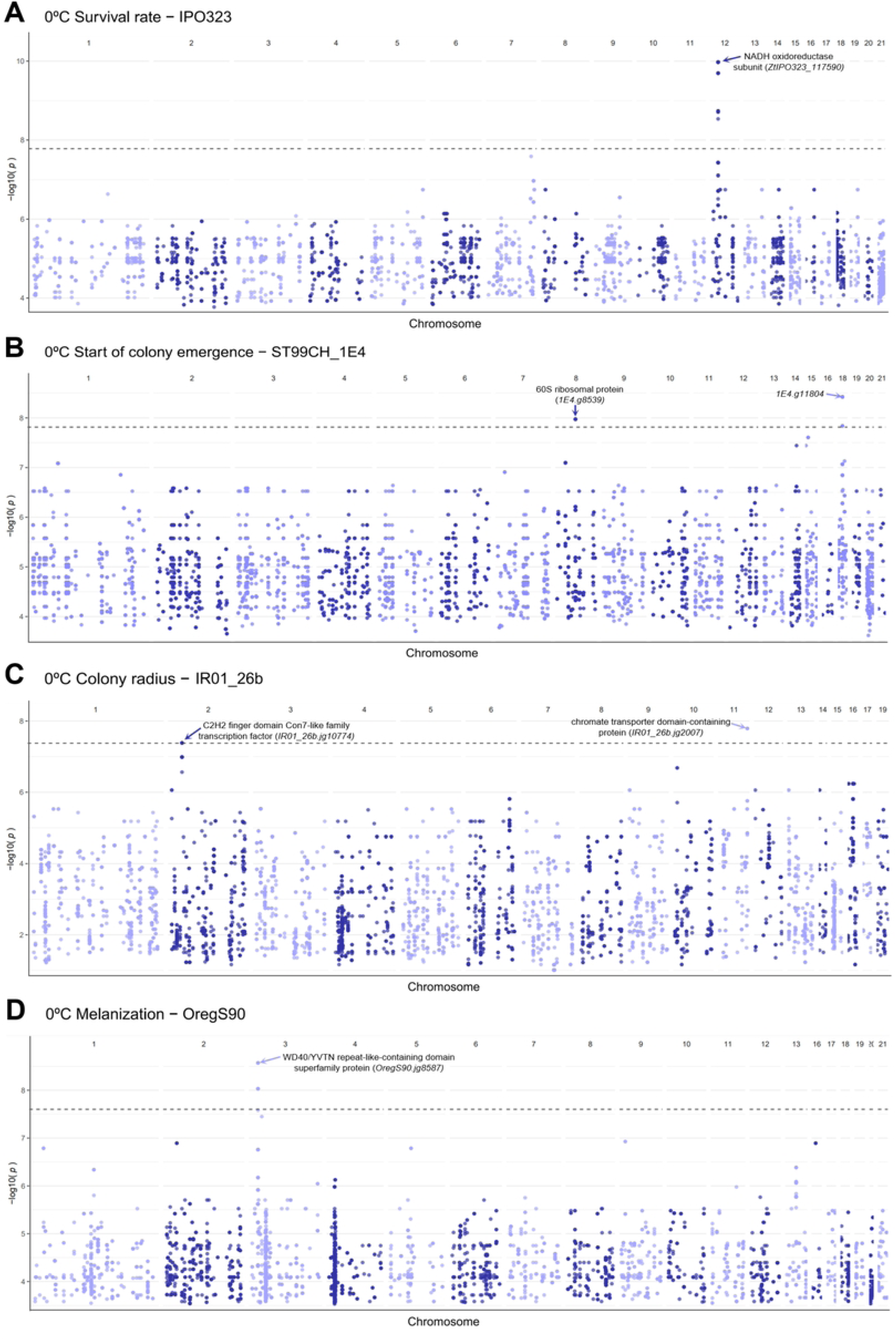
Significant genomic loci and k-mers associated with temperature shock responses in *Zymoseptoria tritici*. The horizontal dashed lines indicate the 10% family- wise error rate significance threshold. Genes co-located with significant k-mers are shown as arrows with corresponding descriptions. (**A**) Manhattan plot of the k-mer GWAS results for the survival rate after cold shock, aligned with the IPO323 reference. (**B**) Manhattan plot of the k-mer GWAS results for the start of colony emergence after cold shock, aligned with the ST99CH_1E4 reference. (**C**) Manhattan plot of the k-mer GWAS results for the start of colony radius 15 days post inoculation after cold shock, aligned with the IR01_26b reference. (**D**) Manhattan plot of the k-mer GWAS results for the melanization 19 days post inoculation after cold shock, aligned with the OregS90 reference.

Next, we assessed the differential expression of the genes co-located with these significant loci in the IPO323 reference. The NADH oxidoreductase 19.3 kDa subunit (ZtIPO323_117590) was not differentially expressed (adj. *p* > 0.05; S7 Table). The 60S ribosomal protein gene (ZtIPO323_092530) was slightly upregulated after cold shock and recovery (log2-fold change = 1.2, adj. *p* < 0.05; S7 Table). The chromate transporter domain-containing protein (ZtIPO323_116130) was also not differentially expressed (adj. *p* > 0.05; S7 Table). The WD40/YVTN repeat-like-containing domain superfamily protein gene (ZtIPO323_038900) was slightly downregulated after heat shock and recovery (log2-fold change = -1, adj. *p* < 0.05; S7 Table). The combination of GWAS and expression datasets showed that about half of the genes significantly associated with post-cold shock phenotypes in the GWAS are differentially expressed in the reference strain IPO323.

## Discussion

Climate change increases the frequency of extreme temperature events, posing challenges for plant pathogen survival. This study investigated how the wheat pathogen *Z. tritici* responds to short-term heat and cold shocks. Using survival, growth, and melanization traits, we assessed *in vitro* phenotypic changes across 122 strains from worldwide geographical origins. Our phenotypic analysis revealed that only the survival- related traits, measured immediately after the shocks, were significantly affected by temperature stress. We performed transcriptomic profiling on the reference strain IPO323 following the temperature shocks and a subsequent recovery period. Gene expression analyses showed distinct transcriptional responses to heat and cold shocks, which stabilized during recovery. Using a multi-reference k-mer-based GWAS, we identified six significant loci associated with four phenotypic traits after cold shock. Our findings highlight specific genetic loci associated with *Z. tritici* phenotypic responses to temperature-induced stress, providing a foundation for future investigations into the genetic basis of resilience to thermal stress in fungal pathogens.

Heat and cold shocks markedly reduced the spore concentration and survival of *Z. tritici* strains immediately following exposure, corroborating prior observations in fungal pathogens [24,38]. However, traits such as colony radius, melanization, and growth rate were resilient, showing no significant deviations from the control condition after a recovery phase. A similar ability to recover after thermal stress was documented in other plant- associated fungi, such as ectomycorrhizal fungi [39]. In addition, our phenotyping analysis revealed a weak but significant correlation between heat and cold shock sensitivities across strains. While most strains exhibited comparable resistance to both types of stress, a minority displayed specialization for either heat or cold stress. This pattern aligns with ecological theories predicting the prevalence of generalist strategies in environments with high variability, where adaptability to a wide range of conditions is advantageous [40,41]. Such generalist strategies have been studied in organisms like insects and amphibians [42–44], and our findings extend this adaptive framework to fungal plant pathogens.

Heat and cold shocks elicit distinct transcriptional responses in *Z. tritici*, illustrating its stress-specific strategies to counter temperature-induced stress. Our analysis reveals a marked reprogramming of the global transcriptome, underscoring differences in gene expression profiles between the heat shock (30°C) and cold shock (0°C) treatments and between immediate shock and recovery phases. Such divergence implies specific molecular pathways are activated under different stress conditions. Excluding the genes of unknown functions, heat shock primarily induced the upregulation of genes associated with protein stabilization and repair, including those involved in posttranslational modifications, protein turnover, and chaperone activity, a phenomenon well-documented in fungal stress responses [45,46]. The induction of genes with functions involved in “posttranslational modification, protein turnover and chaperones” in response to elevated temperatures aligns with their established role in mitigating protein denaturation and aggregation [46,47]. Our detailed investigation of annotated HSPs revealed that most are not differentially expressed after 15h at 30°C. This could be explained by the fine-tuned activation of HSPs at elevated temperatures reported in other systems [48]. The HSP98 and HSP20 proteins were unexpectedly downregulated after temperature shocks. The roles of HSP proteins in stress tolerance, documented in other fungi and organisms [49,50], suggest potential species-specific or context-dependent functions in *Z. tritici*.

Additionally, our data suggest an upregulation of genes involved in secondary metabolite biosynthesis under heat stress, potentially indicating the activation of protective secondary metabolites, as observed in other fungal species in response to abiotic stresses [51]. This is consistent with our observed reduction in spore viability following heat treatment, likely reflecting the cellular cost of maintaining homeostasis and activating protective pathways. Conversely, cold shock in *Z. tritici* led to a widespread downregulation of gene expression, reflecting a cellular strategy to conserve energy and resources under reduced metabolic demand. This response aligns with findings in *Neurospora crassa* and other fungi exposed to cold stress [52,53]. Our findings contribute to a broader understanding of fungal stress biology, highlighting the complex interplay between transcriptional regulation and cellular survival mechanisms under thermal stress. Further exploration into the functional roles of DEGs with unknown functions could unveil novel pathways that underlie fungal resilience to temperature stress, as emphasized by transcriptomic studies in other fungal pathogens [53,54].

Our findings indicate that the transcriptional responses to temperature stress in *Z. tritici* are transient, dissipating rapidly once the stress is alleviated. This explains why exposure to temperature shocks did not significantly impact colony size or morphology. The rapid shift of gene expression suggests efficient recovery mechanisms that restore cellular homeostasis, consistent with reports in other fungal systems where stress-induced transcriptional changes are tightly regulated and short-lived [55,56]. While some DEGs persist after recovery, the limited overlap between heat and cold shock recovery treatments suggests that recovery mechanisms are tailored to the specific type of stress encountered. This stress-specific nature of recovery mechanisms highlights the complexity of fungal adaptation to fluctuating thermal environments. Notably, our phenotypic data on 122 *Z. tritici* strains identified a high variation in phenotypes among strains from the same wheat field in a global sample of fields, with some strains showing higher tolerance to temperature stress than others. Such variation may reflect underlying diversity in genes regulating stress response pathways, suggesting that *Z. tritici* populations likely harbor strain-specific responses to environmental stress conditions. Recent studies in other fungal systems and *Z. tritici* have demonstrated the power of integrating transcriptomics with phenomics to uncover strain-specific adaptations [38,57]. Investigating the transcriptional responses of diverse strains under heat and cold stress conditions could provide deeper insights into the molecular underpinnings of thermal stress resilience in this pathogen.

Our multi-reference k-mer GWAS approach identified six loci linked to phenotypic traits under cold shock, providing evidence for strain-specific adaptation to near-freezing temperature stress within *Z. tritici* populations. Among these, two loci emerged as strong candidates for cold adaptation. The locus associated with cold shock survival co-located with a gene encoding an NADH oxidoreductase subunit. Although not differentially expressed in the reference strain IPO323, this function related to redox reactions and energy metabolism suggests a potential role in enhancing tolerance to oxidative stress, a common consequence of temperature fluctuations [58,59]. Similarly, the locus linked to the initiation of colony emergence was associated with a 60S ribosomal protein gene, which showed slight upregulation after cold shock. Given the essential role of ribosomal proteins in protein synthesis and stress recovery [60], this gene is a compelling candidate for cold stress adaptation. Our findings mark these two genes as novel candidates for cold adaptation, as neither were identified in prior genome-association studies examining long-term temperature conditions or bioclimatic variables [32–37]. This novelty highlights the distinct genetic mechanisms underpinning *Z. tritici* responses to short-term temperature shocks, contrasting with previously studied thermal adaptation.

Our GWAS analysis did not identify significant loci associated with heat shock phenotypes after filtering for transposable elements (TEs). TEs are well-documented drivers of genomic variability and stress adaptation in fungal pathogens [60] and other organisms [61,62]. TE insertions can influence gene regulation by altering promoter activity and chromatin structure or introducing regulatory elements [63,64]. In *Z. tritici*, TEs are mobilized under starvation conditions and during the early stages of host infection [65]. TE-mediated variation can affect melanization and multi-drug resistance phenotypes [66,67]. However, interpreting GWAS-identified TEs is inherently challenging due to their repetitive nature and the difficulty of linking their effects to phenotypic outcomes. In addition, the absence of specific heat shock-associated genes in our GWAS results may reflect the polygenic and complex nature of heat stress responses. Our transcriptomic analyses revealed that over 150 genes were upregulated after heat shock, compared to only 57 following cold shock. This suggests that broader and more intricate gene networks are involved in managing heat-induced stress. Such complexity may hinder the ability of GWAS to detect small-effect alleles that contribute cumulatively to phenotypic variation [68]. Moreover, some *Z. tritici* strains can produce heat-tolerant chlamydospores under elevated temperatures [24,25,69], adding another layer of complexity to strain-specific responses that may not be captured by GWAS, supporting the value of combining multi-omic approaches to unravel complex phenotypic traits in fungal pathogens.

## Materials and Methods

### Strain collection and inoculum preparation

A total of 122 *Z. tritici* strains were used in this study, originating from four geographical regions: the United States (Oregon, 44 strains), Israel (24 strains), Australia (25 strains), and Switzerland (29 strains) (S1 Table). Details on strain sampling can be found in Zhan et al. (2005)[70]. Strains were stored in 50% glycerol or on anhydrous silica at −80°C for long-term preservation. For pre-cultures, we inoculated 150 μL of spore suspension into 50 mL of YPD medium (10 g/L yeast extract, 20 g/L bacto peptone, and 20 g/L dextrose) supplemented with kanamycin (50 mg/L). Cultures were incubated at 18°C and shaken at 120 rpm for 7 days in the dark. Seven-day-old pre-cultures were filtered through two layers of gauze and centrifuged at 3000g for 10 minutes. The spore pellets were resuspended in approximately 20 mL of glucose peptone liquid (GPL) medium (14.3 g/L dextrose, 7.1 g/L bacto peptone, 1.4 g/L yeast extract, and 50 mg/L kanamycin). Two 1:100 dilutions were prepared in water, and spore concentration was estimated using the ImageJ Spore-Counting-V9 pipeline (https://github.com/jalassim/SporeCounter).

### Temperature shock treatments and inoculations for colony phenotyping

Spore suspensions were adjusted to a 7.5 × 10⁵ spores/mL concentration in 1 mL GPL medium. Spore suspensions were exposed to temperature shocks at 0°C (cold shock), 30°C (heat shock), and 18°C (control) for 15 hours, with each treatment performed in triplicate per strain. After the 15-hour incubation, spore concentrations were re-assessed to estimate the first variable, “post-shock spore concentration.”

For colony phenotyping, we adapted the procedure described in [71]. Briefly, we re- assessed spore concentrations after 15h at 0°C, 30°C and 18°C and adjusted the concentrations to 2.5 × 10² spores/mL for the control (18°C) and 5 × 10² spores/mL for the heat and cold shocks. We inoculated 100 μL of the adjusted suspensions onto 35 mL Potato Dextrose Agar (PDA, Difco) Petri plates, incubating them at 18°C in the dark at 70% relative humidity. Three plates were inoculated per strain and treatment.

### Imaging and colony detection

Plates were photographed at 8, 12, 15, and 19 days post-inoculation (dpi) using a Canon EOS 60D camera with the following settings: aperture f/8.0, shutter speed 1/60, and ISO 250. A Sigma 50mm f/2.8 DG Macro lens was used, with manual focus adjusted for each plate. The camera was mounted on a stand at a height of 35 inches (88.9 cm), and images were captured using a remote release to reduce vibration. Images were analyzed using the ImageJ macro “Colony-Analysis-V13” (https://github.com/jalassim/Colony_Analysis), which performed white balancing, colony detection, area measurement, and extraction of per-pixel gray values (mean grayscale values 0-255) for each detected colony. The ImageJ macro settings we used are reported in S1 File. Errors in colony detection were further curated as follows: a subset of 917 randomly selected images from the total 4,392 images was manually curated to classify detected colonies as actual colonies, water shading errors (misidentification of adjacent colonies), or areas incorrectly detected as colonies (S10 Table). This manual classification established a threshold through Principal Component Analysis (PCA) that retained as many actual colonies as possible. Because the dpi influenced the PCA, an individual threshold was set for each time point (S10-S11 Figs).

### Post-temperature shock phenotypic variables

We obtained seven post-shock phenotypic variables, including post-temperature shock spore concentration, survival rate, colony radius, melanization, melanization heterogeneity, growth rate, and start of colony emergence. Figure S12 provides a graphical description of the post-temperature shock phenotypic variables (S12 Fig). We measured the post-temperature shock spore concentration directly by assessing the spore concentration 15 hours after the temperature shock. To estimate the survival rate, we divided the observed number of colonies on the plate by the expected number per plate based on count of inoculated spores post-temperature shock.

We calculated the colony radius using the formula for the area of a circle, following previous studies [71,72]. The image analysis macro provided the colony melanization values by determining the mean grayscale value (scale 0–255; 0 = black, 255 = white) of the pixels within each colony’s area. The melanization heterogeneity depicts the intra-colony melanization variation and is represented by the standard deviation of the grayscale values for pixels within the colony area. We measured the colony radius, melanization, and melanization heterogeneity for all colonies photographed per strain and treatment at 8, 12, 15, and 19 dpi. We performed a linear regression analysis of the radius over time to determine the growth rate and the start of colony emergence. The growth rate corresponds to the slope of the regression line, while the intercept on the *x*-axis indicates the start of colony emergence (i.e., the growth lag phase).

Additionally, we performed Pearson’s correlations between growth variables and treatments to observe their relationships (S13 Fig.; S11 Table). Of the 1,128 comparisons, 651 (58%) showed significant correlations, with absolute correlation coefficients ranging from 0.17 to 0.97 (Pearson’s correlation, *p* < 0.05; S11 Table). Measures of central tendency and other summary statistics for each phenotypic variable per temperature treatment are found in S12 Table. These statistics per geographic population are found in S13 Table.

Finally, we calculated post-temperature shock sensitivity as the ratio between the post- shock and survival rates at the control temperature. We classified the strains based on their post-temperature shock sensitivity scores at 0°C and 30°C. First, we estimated both conditions’ quartiles (0-4) of the post-temperature shock sensitivity. We then classified the strains as belonging to a quartile value for 0°C and for 30°C. For example, a strain with a quartile value of 0 at 0°C has a higher sensitivity to the cold shock, and a strain with a quartile value of 4 at 0°C has a lower sensitivity to the cold shock. We subtracted the quartile value at 0°C from the quartile value at 30°C to obtain a scale of the resistance between cold and heat shocks. The scale was defined as follows: -3 indicates that strains are highly resistant to cold shock compared to heat shock; -2, moderately resistant to cold shock compared to heat shock; -1, slightly resistant to cold shock compared to heat shock; 0, equally sensitive to both heat and cold shocks; 1, slightly resistant to heat shock compared to cold shock; 2, moderately resistant to heat shock compared to cold shock; and 3, highly resistant to heat shock compared to cold shock. S6 Fig provides a diagram illustrating this calculation, and S5 Table contains the corresponding estimates.

### Statistical analyses

We analyzed the phenotypic data using R (v 4.2.1). We used *lmerTest* [73] to perform linear mixed-effects models (LMERs) with the *lmer* function for each phenotypic variable. The LMERs included interaction terms between temperature treatments, strains’ geographic origin, and time points, when applicable. We accounted for strain and experimental batch as random effects. The per colony phenotypic variables were summarized by obtaining the mean per Petri plate to be used as model inputs, as colonies on the same plate were not independent. To investigate fixed effects, we conducted a type III analysis of variance (ANOVA) with the Satterthwaite method. We obtained least-square mean differences through pairwise *t*-tests with the Tukey correction using the *emmeans* package [74].

### RNA sequencing and transcriptome profiling

The pre-culture of the reference strain IPO323 [75] was prepared as described previously for heat and cold shock response phenotyping. We then investigated the transcriptomic response of IPO323 under two experimental conditions: i) A “shock” treatment, where 25 mL spore suspensions (7.5 x 10⁵ spores/mL) were incubated for 15 hours at either 30°C (heat shock), 0°C (cold shock), or 18°C (control), with three replicates for each condition. ii) A “shock and recovery” treatment, where 25 mL spore suspensions were exposed to 15 hours at 0°C, 30°C, or 18°C, followed by an additional 75 hours at 18°C, with three replicates per condition. For each experiment, we transferred 2 mL of spore suspension, centrifuged the samples at approximately 9400g for 2 minutes to remove the media, and immediately flash-froze the spore pellets before storing them at -80°C. The frozen samples were sent for RNA extraction and sequencing to Biomarker Technologies GmbH (BMKgene, Münster, Germany). The sequencing was performed with Illumina NovaSeq X, pair-end 150bp reads. RNA sequencing data is publicly available at Gene Expression Omnibus accession GSE284056.

The sequencing adapters and the reads shorter than 50 bp were trimmed using Trimmomatic v.0.38 [76]: ILLUMINACLIP:${adapters}:2:30:10 LEADING:15 TRAILING:15 SLIDINGWINDOW:5:15 MINLEN:50. The trimmed reads were mapped to the *Z. tritici* IPO323 transcriptome reference [77] and the transcript abundances were quantified using Kallisto v0.46.1 [78] with default parameters for pair-end reads. Read counts were summarized using *tximport* v1.32.0 [79]. The differential gene expression analysis was performed with the R package *DESeq2* [80]. A principal component analysis was performed on the *DESeq2* r-log transformed normalized counts. Genes were considered differentially expressed compared to the controls if the adjusted *p* was less than 0.05 and the log2fold change was higher than 2.5 (upregulated) or lower than -2.5 (downregulated).

Gene Ontology (GO) enrichment analyses were performed for the identified DEGs to assess functional categories. The gene ontology (GO) enrichment analysis was conducted with the R package topGO [81] with the GO annotations obtained by submitting the coding sequence data from the IPO323 reference [77] to the online platform of eggNOG-mapper v2 [82].

### Genome-wide association studies

The genomic sequences of all the strains used were publicly available and the NCBI Bioproject numbers are provided in Supplementary Table S1. We performed a k-mer GWAS as described in [83] as this method increases the power of genome-wide association detection [84]. Briefly, we first performed read trimming using Trimmomatic v0.35 [76], removing the sequencing adapter and low-quality bases with the following parameters: LEADING:15 TRAILING:15 SLIDINGWINDOW:5:15 MINLEN:50. We counted the number of 31bp k-mers for each sample, and we created a k-mer presence/absence table for the 122 strain collection. Following a previously described pipeline [83], we generated a kinship matrix based on the k-mer table to include in a GWAS performed using GEMMA [85] for the seven post-temperature shock phenotyping variables. We assessed the significance of the k-mers using a likelihood ratio test. We considered a k-mer significant if its *p* exceeded the 10% family-wise error rate threshold based on phenotype permutations [83]. This yielded a set of significant k-mers from each association, including redundant k-mer sequences. To remove redundancy, we compiled a list of unique k-mers based solely on their sequence rather than their identifiers, creating a streamlined dataset representing only sequence-unique k-mers across GWAS analyses. To refine the dataset further, we filtered out k-mers found in control conditions. This filtering allowed us to isolate k-mers associated specifically with temperature shock conditions while retaining the control-associated k-mers separately.

One of the advantages of kmer-based GWAS is its non-reference-based approach, allowing for the detection of regions or variants not present in a single reference genome. However, reliance on reference genomes is often necessary to visualize and interpret the results effectively. To overcome this limitation, we constructed a composite reference genome file incorporating the 19 fully assembled *Z. tritici* genomes [31]. We then performed BLAST searches of the k-mer sequences against this composite reference to identify the genomic locations of significant k-mers (https://github.com/afeurtey/Kmer_GWAS/tree/main). We concatenated the mapping of significant k-mers per reference into a single dataset per phenotype, keeping the uniquely mapped k-mers per reference (i.e. avoiding repeats). We then identified “significant k-mer loci” per chromosome as windows containing more than two significant k-mers. If we detected a gap of more than 10kb between the start position of the first significant k-mer and the end position of the last significant k-mers per chromosome, we attributed a new window. We used bedtools v2.31.0 *closest* function with the -d option [86] to identify each reference’s closest neighboring gene or overlapping gene. Each significant k-mer locus was cross-referenced with orthogroups to identify shared and specific loci. Orthology analysis was performed on proteomes from the 19 reference annotations using ProteinOrtho v6.3.2 [87]. Overlapping regions that contained k-mers with common orthogroups or with common k-mers were classified as common significant k-mer loci. Each locus was manually inspected to filter any remaining repeats. To explore the functions of the genes co-located with the significant loci, we referred to the latest annotation of the IPO323 reference [77]. We performed InterProScan searches with the predicted proteins to search for additional predicted functions [88]. Overlap with predicted transposable elements (TEs) was assessed with the latest TEs annotations for the 19 reference genomes for *Z. tritici* [89].

## AI language model assistance

We utilized ChatGPT (developed by OpenAI) to refine the written content of this study. The model offered suggestions and corrections to improve the clarity and grammar of the text based on the authors’ input. All outputs were thoroughly reviewed and revised by the authors to ensure accuracy and alignment with the intended message.

## Author contributions

SMP, AF and CL wrote the manuscript. BAM and JA revised the manuscript. SMP, AF, and CL designed the experiments and analyzed and interpreted the data. SMP acquired the data. JA generated the image analysis tool. SMP, AF, and CL contributed to the conception of the work.

## Acknowledgments

We thank ETH Zurich student Manuel Cavigelli for assistance with preparing phenotyping data. Growth phenotyping data were generated in collaboration with the Genetic Diversity Centre (GDC), ETH Zurich. This project was funded by ETH Zurich. Cecile Lorrain is funded by an SNSF Amizione grant (PZ00P3_209022).

## Supporting information

**S1 Fig. Comparative analysis of colony radius, melanization, and melanization heterogeneity traits across temperature treatments and timepoints.** Boxplots showing colony radius, melanization, and melanization heterogeneity traits measured after 15-hour temperature shock treatments on days post-inoculation not included in Figure 1A. Triangles within each box represent the mean values. Treatments are defined as follows: 0°C for cold shock, 30°C for heat shock, and 18°C as the control. Lowercase letters indicate significant differences between treatments based on linear mixed-effects models per variable and Tukey’s post-hoc pairwise comparisons. *dpi*: days post-inoculation.

S2 Fig. Spore concentration across geographically distinct *Zymoseptoria tritici* populations following temperature shock and control treatments. Boxplots of spore concentration across populations of different geographical origins after a 15-hour temperature shock or control treatment. Triangles within each box indicate mean values. Treatments are defined as follows: 0°C for cold shock, 30°C for heat shock, and 18°C as the control. Arrows and asterisks denote significant differences between treatments and populations, as determined by a linear mixed-effects model with Tukey’s post-hoc pairwise comparisons (*: *p* < 0.05; **: *p* < 0.01; ***: *p* < 0.001).

**S3 Fig. Survival rates across *Zymoseptoria tritici* populations after exposure to temperature shocks and control conditions.** Boxplots of survival rate across populations of different geographical origins after a 15-hour temperature shock or control treatment. Triangles within each box indicate mean values. Treatments are defined as follows: 0°C for cold shock, 30°C for heat shock, and 18°C as the control. Arrows and asterisks denote significant differences between treatments and populations, as determined by a linear mixed-effects model with Tukey’s post-hoc pairwise comparisons (*: *p* < 0.05; **: *p* < 0.01; ***: *p* < 0.001).

**S4 Fig. Survival-based sensitivity to temperature shocks across populations.** Boxplots of the survival-based sensitivity to temperature shocks compared to the control for each population. The dashed line at “1” indicates equal survival rates between treatments and the control. There are no significant differences between treatments and populations based on the linear mixed-effects model and Tukey’s post-hoc pairwise comparisons.

**S5 Fig. Correlation analysis of population-level sensitivity to heat and cold shock treatments.** Scatter plots of survival-based sensitivity following heat shock (30°C) versus cold shock (0°C) per population. Pearson’s correlation coefficient (*R*) and *p* values are displayed at the top of the plot.

**S6 Fig. Visual representation of the calculations of the relative resistance to temperature shocks.**

**S7 Fig. GO categories of DEGs in *Zymoseptoria tritici* under temperature shock treatments.** A) Proportion of DEGs with GO annotations under cold shock treatment. B) Detailed categories of DEGs under cold shock treatment. C) Proportion of DEGs with GO annotations under heat shock treatment. D) Detailed categories of DEGs under heat shock treatment.

S8 Fig. Heatmap depicting expression levels of predicted chaperone heat-shock proteins in *Zymoseptoria tritici* under temperature shock and control conditions. Samples were collected either immediately after treatment (15 h) or following a recovery period (15 h at treatment + 75 h at 18°C). Rows represent individual genes, and columns indicate samples from three conditions: cold shock (0°C), heat shock (30°C), and control (18°C). Color gradients denote relative expression levels. Genes were grouped by hierarchical clustering based on similar expression patterns across treatments.

**S9 Fig. GO categories of DEGs in *Zymoseptoria tritici* under temperature shock-and-recovery treatments.** A) Proportion of DEGs with GO annotations under cold shock- and-recovery treatment. B) Detailed categories of DEGs under cold shock-and-recovery treatment. C) Proportion of DEGs with GO annotations under heat shock-and-recovery treatment. D) Detailed categories of DEGs under heat shock-and-recovery treatment.

**S10 Fig. Principal component analysis (PCA) of phenotypic data across days post-inoculation, highlighting manually curated colony recognition errors.** PCA plots show colony phenotypic data for 122 *Zymoseptoria tritici* strains across different days post-inoculation (dpi) under temperature shock and control treatments. The analysis is based on seven phenotypic traits. Colors represent distinct types of colony recognition errors identified during manual curation, with colonies not curated manually shown in gray. Black lines mark cut-off thresholds used to filter colony recognition errors for each dpi. Three categories of colony recognition errors are highlighted: *Background*, referring to areas incorrectly identified as colonies; *Real_Colony*, representing colonies correctly detected; and *Watershedding*, indicating errors caused by the misidentification of adjacent colonies.

**S11 Fig. Proportion of colonies retained based on PCA filtering, shown by manually curated data categories and unseen data across days post-inoculation.** This figure presents the proportion of colonies from 122 *Zymoseptoria tritici* strains retained for further analysis after PCA-based filtering, as determined by manually curated data categories and unseen data at different days post-inoculation. Colonies were categorized into four groups: *Background*, representing areas incorrectly detected as colonies; *Real_Colony*, referring to colonies correctly identified; *Watershedding*, indicating errors caused by the misidentification of adjacent colonies; and *unknown*, corresponding to unseen data that were not manually curated.

S12 Fig. Methodological overview for quantifying seven phenotypic variables in *Zymoseptoria tritici* colonies post temperature shock. Visual summary of the methods used to derive seven phenotypic variables: survival rate, colony emergence timing, growth rate, colony radius, melanization, and melanization heterogeneity.

**S13 Fig. Correlation plot of the seven phenotypic variables across temperature shock and control treatments.** Colors represent Pearson’s correlation coefficients, while asterisks indicate significance levels (*: *p* < 0.05; **: *p* < 0.01; ***: *p* < 0.001).

S1 Table. Comprehensive metadata of *Zymoseptoria tritici* strains, including origin, sampling year, and sequencing availability. Detailed information for each strain, including its name, genetic cluster assignment, geographical location, inferred coordinates of the sampling site, sampling year, and associated BioProject for sequencing data.

**S2 Table**. **Analysis of Variance (ANOVA) results for Linear Mixed Effects Models (LMER) of the seven phenotypic variables after treatment.**

**S3 Table. Pairwise Tukey post hoc test results for Linear Mixed Effects Model (LMER) of the post-temperature shock spore concentration per treatment and per treatment and geographical origin.**

**S4 Table. Pairwise Tukey post hoc test results for Linear Mixed Effects Model (LMER) of the post-temperature shock survival rate per treatment and per treatment and geographical origin.**

**S5 Table. Per strain relative resistance to temperature shocks.** The description of the categories can be found in the Materials and Methods section and S6 Fig.

**S6 Table. Count and percentage of the strains per category of relative resistance to temperature shocks.** The explanation of the categories can be found in the Materials and Methods section and S6 Fig.

**S7 Table. Differential gene expression results from temperature shocks at two timepoints.** For the “Condition” columns, *S* represents shock treatment, and *R* represents the shock-and-recovery treatment.

**S8 Table. All k-mers significantly associated with each of the 48 phenotypes.** This includes variables under the temperature shocks and the control.

**S9 Table. Significant genomic loci associated with heat and cold shock treatment phenotypes.**

**S10 Table. Curated subset of images with annotated colony identifications, categorized as true colonies or artifacts.**

S11 Table. Correlations between the seven phenotypic variables per treatment. Both Pearson’s *R* and *p* values are presented.

**S12 Table. Summary statistics of phenotypic variables per treatment.**

**S13 Table. Summary statistics of phenotypic variables, per treatment and geographic origin.**

**S1 Data. Source data for phenotypic analyses.**

**S1 File. Settings ImageJ Macro Colony Analysis**

